# Tissue Dynamics Spectroscopic Imaging: Functional Imaging of Heterogeneous Cancer Tissue

**DOI:** 10.1101/2020.09.24.295337

**Authors:** Zhe Li, Bihe Hu, Guang Li, Sharon E. Fox, Shadia Jalal, John Turek, J. Quincy Brown, David D. Nolte

**Author notes:** Address all correspondence to: David D. Nolte.

## Abstract

**Significance:** Tumor heterogeneity poses a challenge for the chemotherapeutic treatment of cancer. Tissue dynamics spectroscopy (TDS) captures dynamic contrast and can capture the response of living tissue to applied therapeutics, but the current analysis averages over the complicated spatial response of living biopsy samples.

**Aim:** To develop tissue dynamics spectroscopic imaging (TDSI) to map the heterogeneous spatial response of tumor tissue to anticancer drugs.

**Approach:** TDSI is applied to tumor spheroids grown from cell lines and to *ex vivo* living esophageal biopsy samples. Doppler fluctuation spectroscopy is performed on a voxel basis to extract spatial maps of biodynamic biomarkers. Functional images and bivariate spatial maps are produced using a bivariate color merge to represent the spatial distribution of pairs of signed drug-response biodynamic biomarkers.

**Results:** We have mapped the spatial variability of drug responses within biopsies and have tracked sample-to-sample variability. Sample heterogeneity observed in the biodynamic maps is associated with histological heterogeneity observed using inverted Selective-Plane Illumination Microscopy (iSPIM).

**Conclusion:** We have demonstrated the utility of TDSI as a functional imaging method to measure tumor heterogeneity and its potential for use in drug-response profiling.

## 1. Introduction

Tumor heterogeneity presents a challenge for the successful treatment of cancer using chemotherapeutics(1). For instance, genetic variability in tumors caused by clonal outgrowth of selected genotypes within a tumor may cause subsets of cells with genetic variations to be resistant even while the majority of the tumor responds to treatment. Selective pressure and genetic drift of the cancer cell population during treatment often leads to patient relapse and the emergence of broad chemoresistance and refractory disease(2-4). In addition to genetic heterogeneity, there is also spatial heterogeneity in tumor tissue arising from varying tissue constituents as well as varying microenvironments, including differences in extracellular matrix and connective tissues. The tumor microenvironment(5, 6) and epigenetic variations(5, 7-9) pose significant challenges to the selection of treatment based on genetic profiles. This has led, as an alternative, to phenotypic profiling(10-12) of cancer tissue that captures the systemic response of cancer tissue to applied therapy. The challenge for phenotypic profiling of cancer tissue is the need to image intact microenvironments deep inside tissue, far from surface damage caused by surgical resection, and deep inside transport-limited regions that experience hypoxia, nutrient depletion and metabolite build-up.

Optical coherence imaging (OCI)(13, 14) is a deep-tissue coherence-domain imaging approach based on digital holography(15-17) that is a form of full-frame optical coherence tomography (FF-OCT)(18, 19). Dynamic speckle in OCI images caused by dynamic light scattering from intracellular motions enables biodynamic imaging (BDI)(20) to use intracellular dynamics as a unique form of image contrast. The changes in intracellular motions caused by applied therapeutics have been studied using tissue dynamics spectroscopy (TDS)(21) to separate the effects of drugs across broad spectral bands and to capture specific signatures from different classes of drugs with different mechanisms of action(22). Preclinical trials of therapy responsivity assessment have been completed using TDS in spontaneous canine B-cell lymphoma and in ovarian xenografts(23, 24). In a substantially different setting, assisted reproductive technology (ART) correlates the viability of cumulus-oocyte complexes with parameters from sample fluctuation power spectra(25).

The methodology of TDS is usually applied to entire samples that can be as large as 1 mm^3^ in volume (e.g. biopsies). However, intra-sample variability in the TDS signatures poses a challenge for the prediction of patient response to therapy. While previous work using TDS has identified and characterized the different baseline conditions and drug responses in the “shell” and “core” areas of the samples(22, 25, 26), in that analysis, the boundary between the shell and core was arbitrarily defined. Some samples have a more complicated drug response structure than a simple “shell” and “core” model, as shown in Fig. 1, where the sample shows variation in both strength and pattern in its drug response in the two areas.

**Fig. 1.**
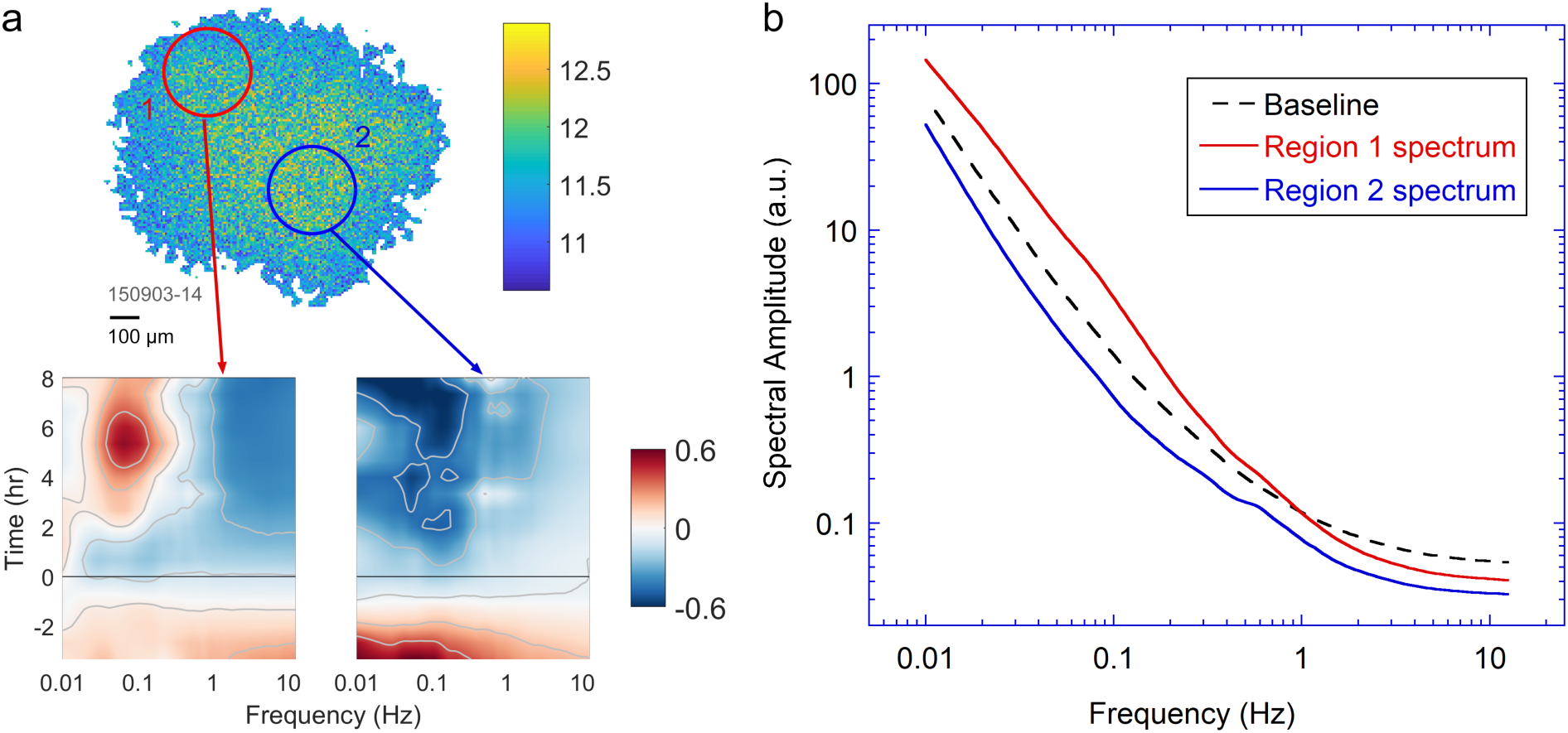
a) An OCI image of a human esophageal biopsy and differential spectrograms (defined below) for the two circled areas. At time *t* = 0 (black line), 100 mL of 25 μM cisplatin and 25 μM fluorouracil (5fu) combined solution is added to the sample. The two regions have significantly different responses: Region 1 shows enhancement in low frequency and suppression in high frequency (“redshift”), while region 2 has suppression across all frequencies. b) The terminal spectra of region 1 and 2, respectively, compared to the average sample baseline spectrum.

To address this problem, we introduce a functional imaging method called tissue dynamics spectroscopic imaging (TDSI) that evaluates sample response on a pixel level. In addition to a full-duration response map, the response can be segmented along the time axis to derive the time-lapse evolution of drug response, which can reveal the different rates at which a drug acts on each area. This methodology offers a quick, intuitive visualization of sample heterogeneity and drug effects. TDSI differs from dynamic-contrast OCT performed with off-axis holography (27), Full-field OCT (28) or conventional OCT (29-34) because its imaging contrast arises from shifts in intracellular dynamics caused by applied therapeutics rather than from baseline intracellular dynamics. In this way, TDSI is functional imaging that is specific to drug efficacy and displays the heterogeneous response of tissue to therapies.

## 2. Materials and Methods

### 2.1 Sample Preparation

Biological samples used in this paper include tumor spheroids grown from the DLD-1 intestinal adenocarcinoma cell line (ATTC, Manassas, VA) and human esophageal tumor biopsies. (IRB-approved IUCRO-0486 clinical trial. Written consent was obtained for each patient.) The multicellular DLD-1 spheroids mimic small avascular *in vivo* tumors by having an active growth zone on the outside of the spheroid and regions of apoptotic and necrotic cells progressively towards the core as nutrients and oxygen become rate-limiting for cell growth, but otherwise the spheroids are highly homogeneous. In contrast, the esophageal tumors are highly heterogeneous, with complex physiology and consist of different cell types. DLD-1 spheroids were grown in Corning U-bottom 96-well spheroid plates, and esophageal tissues were obtained by pinch biopsy from human patients. Tumor biopsies were collected and transported in chilled RPMI-1640 medium with HEPES buffer and cut into small pieces of 1 mm size or less. Both types of samples were immobilized in 1% low-gel-temperature agarose in the RPMI-1640 basal medium. Immobilized samples were overlaid with RPMI-1640 containing 10% heat-inactivated fetal calf serum (Atlanta Biologicals), penicillin (100 IU), and streptomycin (100 μg/mL).

Sample heterogeneity observed using TDSI was verified with high-resolution 3D images obtained from inverted selective plane illumination microscopy (iSPIM). To prepare samples for iSPIM imaging, 10% neutral buffered formalin (NBF) was injected into each well to fix the tissues. After being washed with PBS, the samples were stained with 50 μM DRAQ5 (Biostatus, Ltd.) overnight with gentle agitation, then 0.5 mg/mL 80% ethanol-based Eosin Y (E4009, Sigma-Aldrich) for 4 hours. Afterwards, the agarose embedding the samples in the dish wells was removed, and samples were then washed with DI water three times and PBS once followed by being immersed in X-CLARITY mounting solution (Logos Biosystems) for 15 minutes. Finally, the samples were fixed in the imaging chamber with silicone glue and immersed in X-CLARITY mounting solution for iSPIM imaging. After being imaged, the samples were processed for traditional H&E through the Tulane Medical School Histology Department. Four-micrometer thick sections were cut and stained until each tissue was exhausted. The total processing time is about 16.5h in total after fixation, including the PBS/di-water washing time. Imaging of each sample takes about 1 min average scanning time. Full-resolution image processing of each sample takes approximately 50 minutes, including time for reconstruction, making pseudo-color images, and visualization.

### 2.2 Tissue Dynamics Spectroscopy

Speckle fluctuation dynamics of biological samples were measured and analyzed using a biodynamic imaging system. The system optical configuration is a Mach-Zehnder interferometer, shown in Fig. 2. The light source is a Superlum S840-B-I-20 superluminescent diode with a center wavelength at 836.2 nm and a full power output of 22.9 mW with a bandwidth of 50 nm and a coherence length of approximately 10 μm. A QImaging EMC2 camera is used for image acquisition. Lenses L5 and L6 constitute a 4f imaging system to an image plane (IP). The image plane is subsequently transformed by lens L7 to the Fourier plane located at the CCD pixel array. A single sample image is obtained by performing a 2D FFT on the hologram captured by the CCD camera located on the Fourier domain, and one of the two conjugate images is stored as a 256 × 256 array(26). Because of the use of long focal lengths (f_5_ = f_6_ = 150 mm), the speckle size on the camera plane is approximately 60 μm, and the reconstructed object-plane point-spread function is approximately 15 μm spanned by three pixels.

**Fig. 2.**
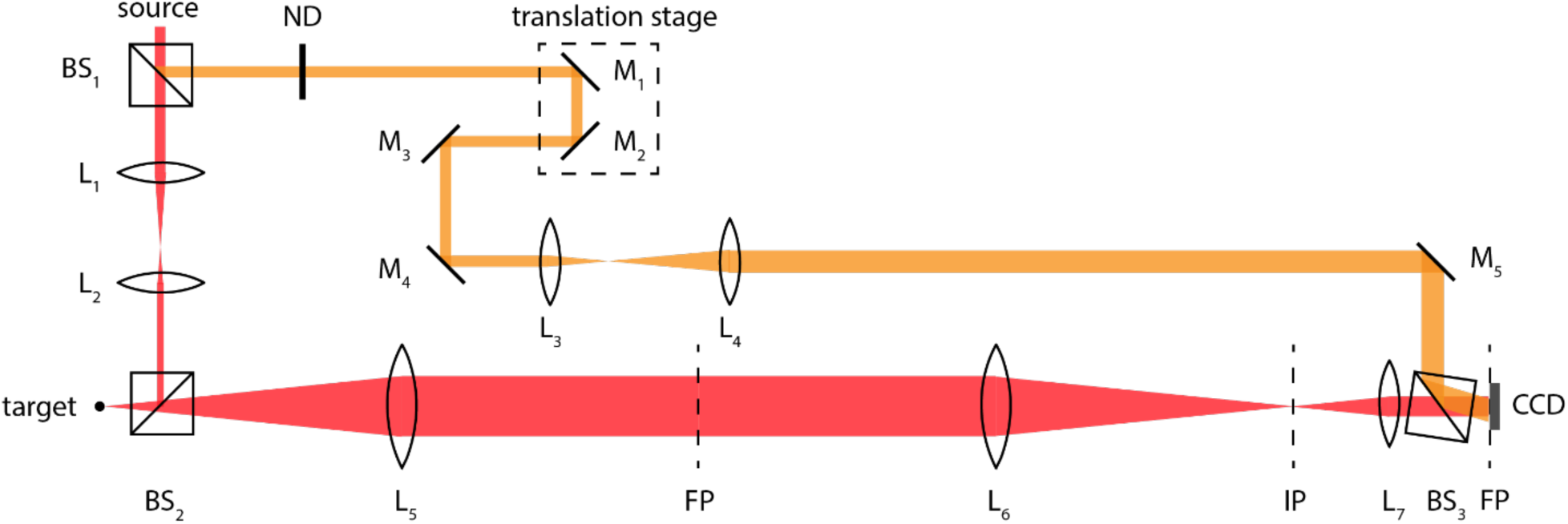
A schematic of the interferometer with a Mach-Zehnder off-axis digital holography configuration. The camera is on the Fourier plane. Optical sections are reconstructed using a 2D spatial Fourier transform. ND: neutral density filter. L_1_ – L_7_: lenses. BS_1_ – BS_3_: beam splitters. M_1_-M_5_: mirrors. FP: Fourier plane. IP: image plane. f_5_ = f_6_ = 150 mm. f_7_ = 50 mm. The translation stage defines the coherence gate for time-domain ranging.

Two data acquisition formats are used for data presented in this paper: A format containing 2048 frames captured at 25 fps, and a format with 500 frames at 25 fps and 50 frames at 0.5 fps. The second format has a smaller data storage requirement but requires a stitching algorithm to create a single spectrum(35). Within this paper, only DLD samples use the 2048 frame format, while the esophageal samples used the 500/50 frame format to assure comparability among spectra within a given study. A typical experiment has 6 time-frames of baseline measurements and 15 time-frames of drug response measurements looping repeatedly through 16 wells in a 96-well plate format. A single loop through all 16 samples takes about 40 minutes.

The fluctuation power spectrum from one pixel at position (*i, j*) in the sample is calculated as the square of the FFT of the intensity time series:

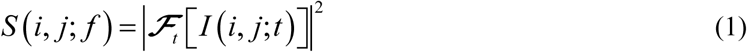

The average spectrum of a region *σ* is calculated as

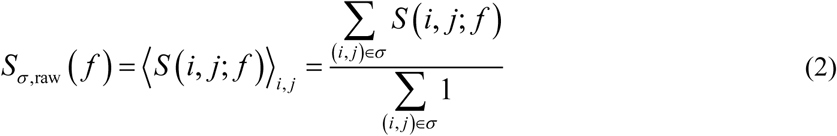

where *σ* is a spatial mask segmenting the entire sample. The “raw” spectrum is normalized based on Parseval’s theorem(35):

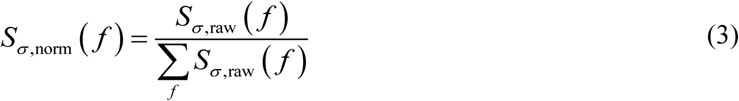

i.e. dividing the raw spectrum by the sum of all frequency components (including the DC component). The full-length spectrogram of segment *σ* is a time-lapse sequence of differential spectra where each spectrum at time *τ* is calculated as a normalized logged spectrum subtracted by the baseline spectrum averaged over the full sample:

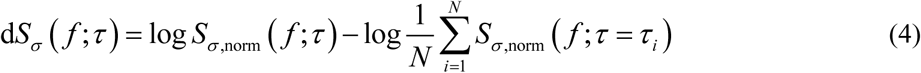

where *N* is the number of baseline loops.

### 2.3 Biodynamic biomarkers and TDS visualization

Biomarkers that evaluate sample preconditions, also called baseline conditions, before a treatment is applied include backscatter brightness (“BB”), normalized standard deviation (“NSD”), spectrum knee frequency (“KNEE”) and <u>spectral slope (“S”) (33)</u>. Biomarkers that evaluate drug responses include changes in these biomarkers after application of drugs as well as features extracted from spectrograms. This paper focuses mainly on the drug response and features from spectrograms.

Despite the differences in drug responses related to sample baseline conditions and drug mechanisms, a biodynamic drug spectrogram usually has one of a limited number of patterns. Spectroscopic masks (time-frequency filters) are designed to match the characteristics of the spectrograms, a few of which are shown in Fig. 3a. The top 3 patterns G0, G1 and G2 form a set of “orthonormal” masks that are related to the broadband (in the sense of frequency components) pattern of a spectrogram, while the bottom 3 patterns FL, FM, and FH form another set of masks related to local response patterns. These frequency bands, and their associated frequency cutoffs, can be related to changes in intracellular motions and their related speeds (23, 36). Although the individual biomarker values depend on the choice of cut-off frequencies, the orthonormal character of the masks provide a unique representation of the spectrogram, and this representation is used in pattern recognition algorithms. For each mask, a feature value is obtained by calculating the inner products of the spectrogram and the mask, i.e. projecting the spectrogram onto the mask (22), and the features of a spectrogram are represented by a vector of feature values.

**Fig. 3.**
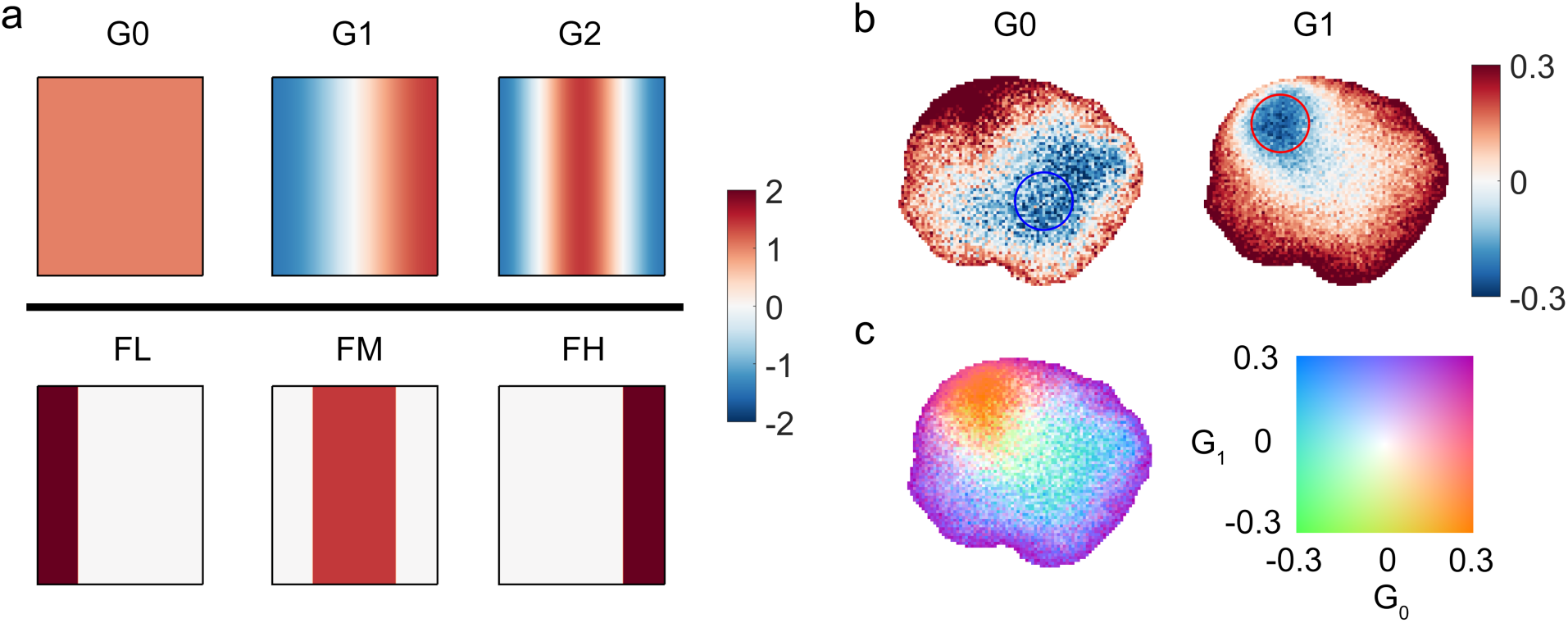
a) A subset of the spectrogram masks used in the color merge. b) Two maps of drug response of an esophageal sample “150903-14” (same as in Fig. 1) exposed to cisplatin and fluorouracil combination therapy under masks G0 and G1. c) A “merged” bivariate color image with its 2D color map.

After condensed-data-format files are generated (See supplement section S.1), a differential spectrogram for each TDSI pixel (referred to as a “microspectrogram”) is calculated as

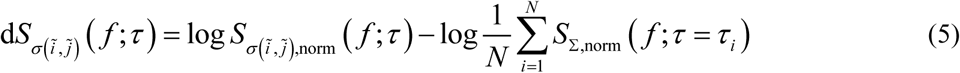

where the average baseline spectrum of the entire sample is used instead of the TDSI pixel’s “own” baseline. For a given mask, a feature value (discussed above) can be calculated for each pixel, and the feature values of an entire sample produce a 2D image called a TDS image (TDSI). Two TDS images under the “G1” mask (blue shift associated with increased intracellular speeds) and “G2” mask (mid-frequency enhancement often associated with active membrane processes) are shown in Fig. 3b. For the “G1” TDS image, the area in the blue circle has negative values, indicating the inverse of the “G1” mask (i.e. the “blue-red” pattern spectral response shown in Fig 3a), which matches the differential spectrogram of the circled area of the same sample as shown in Fig 1a. Similarly, the “blue-red-blue” spectral response in the red circle area agrees with the positive values in the same area in the “G2” TDS image.

As discussed above, the sample in Fig. 3 has a large variation in drug response in terms of strength and spectral patterns, displayed by the distribution of values in the TDS images (TDSI) of Fig. 3b. Both “G0” and “G1” images show a change in magnitude and sign from bottom left to top right (corresponding to changes in the strength of drug response, referred to as “intramask heterogeneity”), and the “G0” image has a different pattern than “G1”, where “G0” has strong negative values on the bottom right while “G1” has strong negative values in the upper left (corresponding to changes in pattern, referred to as “intermask heterogeneity”). To better visualize the variation, bivariate color images are introduced to produce a single visualization that captures drug responses across the entire sample, where each “variable” is a feature value of a mask. Feature values from two masks are a convenient way to illustrate drug-response heterogeneity within a sample, and the following discussion will focus on bivariate representations of drug response.

Bivariate color maps are used in cartography (37, 38) and medical imaging (39), and many studies have addressed how to choose proper colormaps for bivariate data visualization (40-42). We have elected to use the “Teuling3” colormap in the following visualizations, which is generated by linearly interpolating four colors at the four corners in the sRGB space plus a “whitening” core in the center (40, 43). This color map has good color saturation, relatively equal visual impact, and a zero value appears as white, which is consistent with the diverging “blue-red” 1D colormap used in our spectrograms and univariate TDS maps. Fig. 3c shows a bivariate image of a human esophageal biopsy sample “merged” from the two univariate TDS maps in Fig. 3b.

When a sample has areas with spectrograms that are the inverse of each other, they cancel each other out in an average over the full sample, resulting in a mild spectrogram and near-zero feature values. In this case, the weak average response belies the strong change in the intracellular dynamics and the fluctuation spectra, and this can lead to the misinterpretation that the sample does not respond to the drug. To address this problem, two new biomarkers that evaluate sample heterogeneity are added to the “traditional” average spectrogram-based biomarkers. The two biomarkers evaluate the “intramask” and “intermask” heterogeneity respectively. To achieve a high signal-to-noise ratio, TDS images are first (re)generated with an 8 × 8 pixel averaging (instead of the standard 2 × 2 px). The coarse eight-pixel averaging is used only to calculate the heterogeneity biomarker values of a sample at a scale of 40 microns and above. Finer scale is not necessarily meaningful for quantifying spatial heterogeneity across a millimeter sample. The feature values are bounded to a range [−*A*_th_, *A*_th_] and then assigned “scores” ranging from 0 to 1 for both heterogeneity evaluations, calculated using the following steps:

1. Select a set of *n* masks
2. For each mask *u*, calculate Δ*a*^*u*^ and Δ [sgn (*a*^*u*^)]
3. For each mask pair *u*-*v*, calculate | *ρ* (*a*^*u*^, *a*^*v*^) and | | *ρ* [sgn (*a*^*u*^), sgn (*a*^*v*^)]|
4. The first biomarker denoting overall intra-mask heterogeneity is calculated as

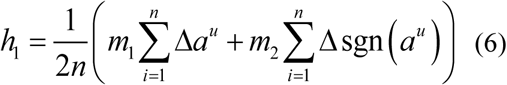
5. And the second biomarker representing overall inter-mask heterogeneity is calculated as

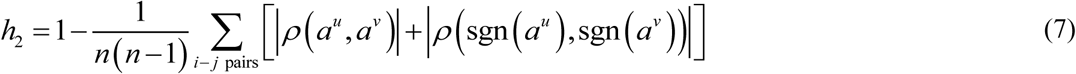

where 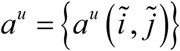 are values of the TDS image of mask *u*, sgn (*a*^*u*^) is a map of signs of *a*^*u*^, Δ*a*^*u*^ is the standard deviation of *a*^*u*^, *ρ* (*a*^*u*^, *a*^*v*^) is the correlation coefficient of *a*^*u*^ and *a*^*v*^, and *m*_1_ and *m*_2_ are normalization factors

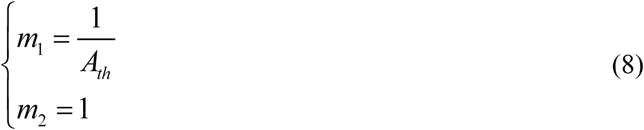

based on Popoviciu’s inequality on variances (44). We use *A*_th_ = 0.3 and *n* = 3, and the local masks FL, FM and FH are used for heterogeneity scores.

Based on these definitions, the extreme values are achieved under these cases: *h*_1_=0 when *n* TDS images are completely uniform (totally homogeneous), and *h*_1_=1 when TDS images have only two values of opposite signs in an equal number of pixels. When the TDS images are completely nonlinearly correlated to each other, then *h*_2_=0. In the opposite case when all values are completely correlated then *h*_2_=1. Examples will be provided in the next section to illustrate these two heterogeneity benchmarks.

### 2.4 Time-lapse drug response visualization

After a drug is added to a biopsy sample, the change in its intracellular dynamics is usually not immediate and depends on the drug mechanism of action, especially for drugs targeting DNA <u>that produce slow cellular responses</u>. Also, some parts of the sample may respond to a drug faster than the entire sample. TDSI allows us to study <u>both the time delay</u> and non-uniformity in drug action, which is called time-lapse TDSI. Instead of extracting feature values from full-length spectrograms, time-lapse TDSI uses responses within a small moving time “window” of the spectrogram. Examples are included in the following sections.

### 2.5 Inverted Selective Plane Illumination Microscopy (iSPIM)

The iSPIM system is constructed around a commercially available diSPIM platform (ASI) and has been described in previous publications(45, 46). In brief, two immersion objectives (CTO, ASI/Special Optics, 15.3X – 17.9X) are orthogonally mounted above the sample, with each at a 45° angle from the norm, enabling traditional sample preparation. Dual-view imaging is possible by alternating roles of the two objectives as illumination and detection, but only single-view was adopted for this paper. Volumetric images were obtained by moving the sample with respect to the objectives. To cover the whole area of the sample, multiple y strips were acquired with about 20% overlap between two adjacent strips. After imaging was completed, the images were first shifted and interpolated with custom MATLAB code to recover its 45° angle(47). Multiple paths were then stitched with Fiji plugin(48). The DRAQ5 and eosin images (D&E) were remapped to a composite RGB stack to simulate the traditional H&E colors(49). Finally, 3D reconstruction of the D&E images was obtained from the alpha blending mode of 3D viewer of Vaa3D (50, 51). The mounting media is Xclarity with a refractive index of **1.45**, that when combined with the **NA of 0.4**, gives a magnification M of 16.7X with a lateral resolution of 0.76 μm and an axial resolution of 3.8 μm.

## 3. TDSI Results

A large number of esophageal biopsies display spatially heterogeneous responses to drugs. InFig. 4a, two biopsy samples that have large intra-mask heterogeneity are presented in univariate and bivariate forms. Sample “151208-6” has a weak response when averaged over the whole-sample spectrogram (Fig. 4b “global”), because the local areas “1” and “2” have strong but opposite responses that tend to cancel in the sample average.

**Fig. 4.**
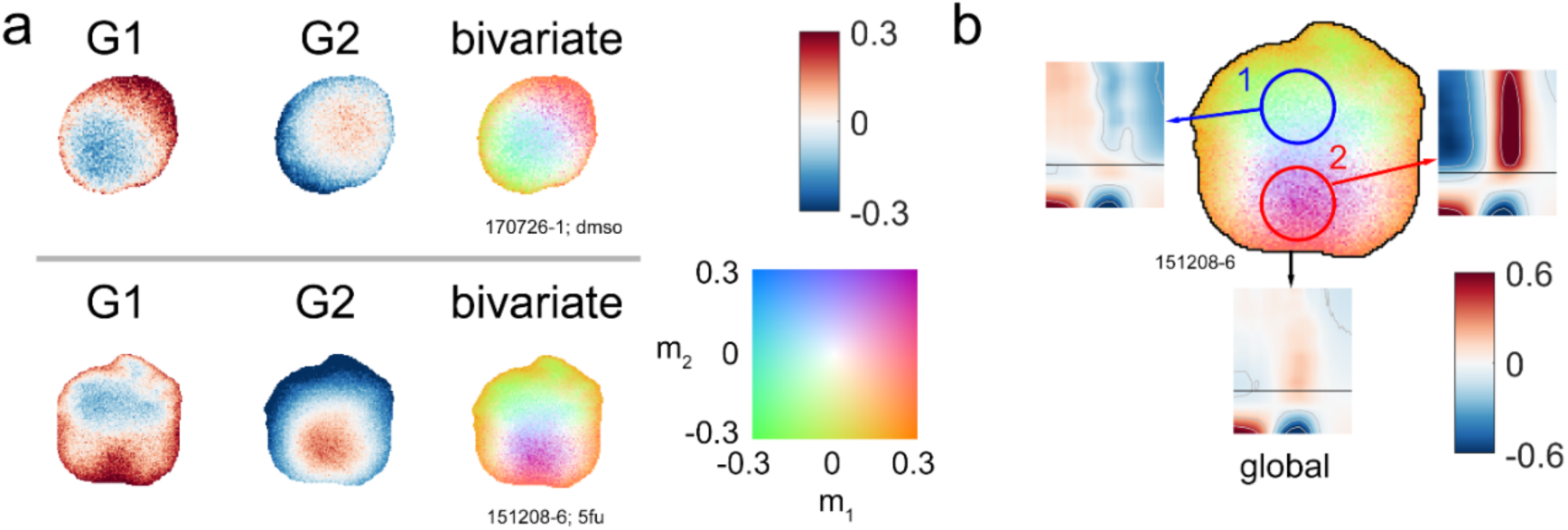
a) Bivariate color representation of drug responses of 2 samples treated with different drugs, showing two univariate maps and a “merged” bivariate color map. The first sample was refreshed with DMSO, while the second sample was treated with 25 μM fluorouracil (5fu). b) Global and regional spectrograms of sample “151208-6” from a). The global spectrogram has a relatively weak response (max<10%), while the two circled areas have 30%-60% enhancement or suppression. Drugs were added at *t*=0 (black line on spectrograms).

An assortment of bivariate TDS images is shown in Fig. 5. Some samples have relatively uniform color in the “merged” map, indicating smaller variation in the biomarker values, while others have a rainbow-like smooth transition across the sample, which is related to high heterogeneity in the drug response.

**Fig. 5.**
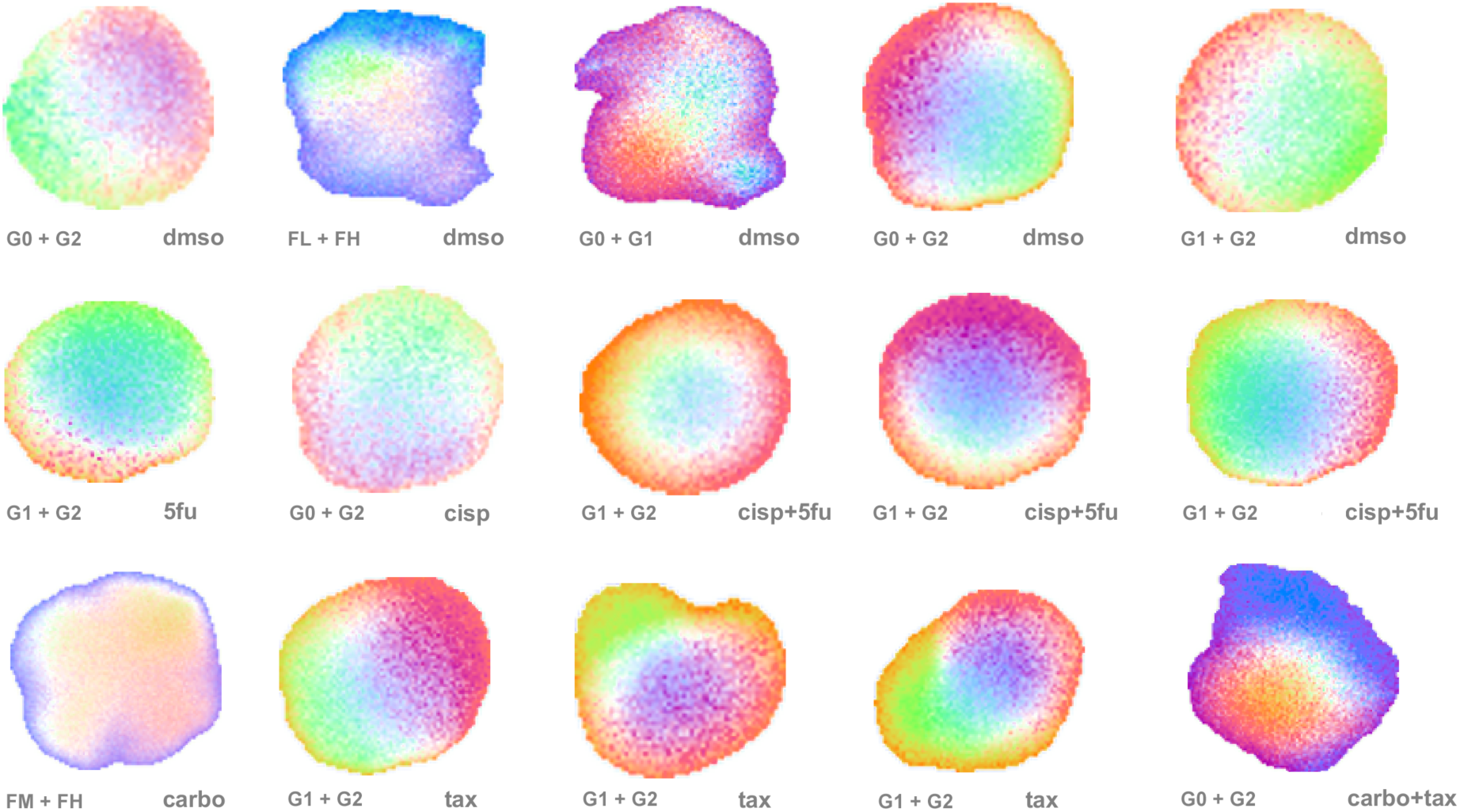
More examples of bivariate TDS images showing sample-to-sample variability in drug responses. Masks are designated in the lower left corners of images, while lower right corners designate drug treatments. Drug abbreviations: <u>dmso: 0.1% DMSO in growth medium (used as a negative control),</u> cisp: 25 μM cisplatin, 5fu: 25 μM fluorouracil, tax: 5 μM paclitaxel, carbo: 25 μM carboplatin. “+” indicates a combination of two drugs. The color map and scale are the same as in Fig. 3.

There are roughly three types of heterogeneity, shown in Fig. 6 along with their *h*_1_ and *h*_2_ scores: (i) Type I are samples that have almost uniform responses under all masks. (ii) Type II samples have spatial variability but show similar patterns across different masks. (iii) Type III samples have TDS images with non-overlapping strongly responding areas. Types I and III are the most common.

**Fig. 6.**
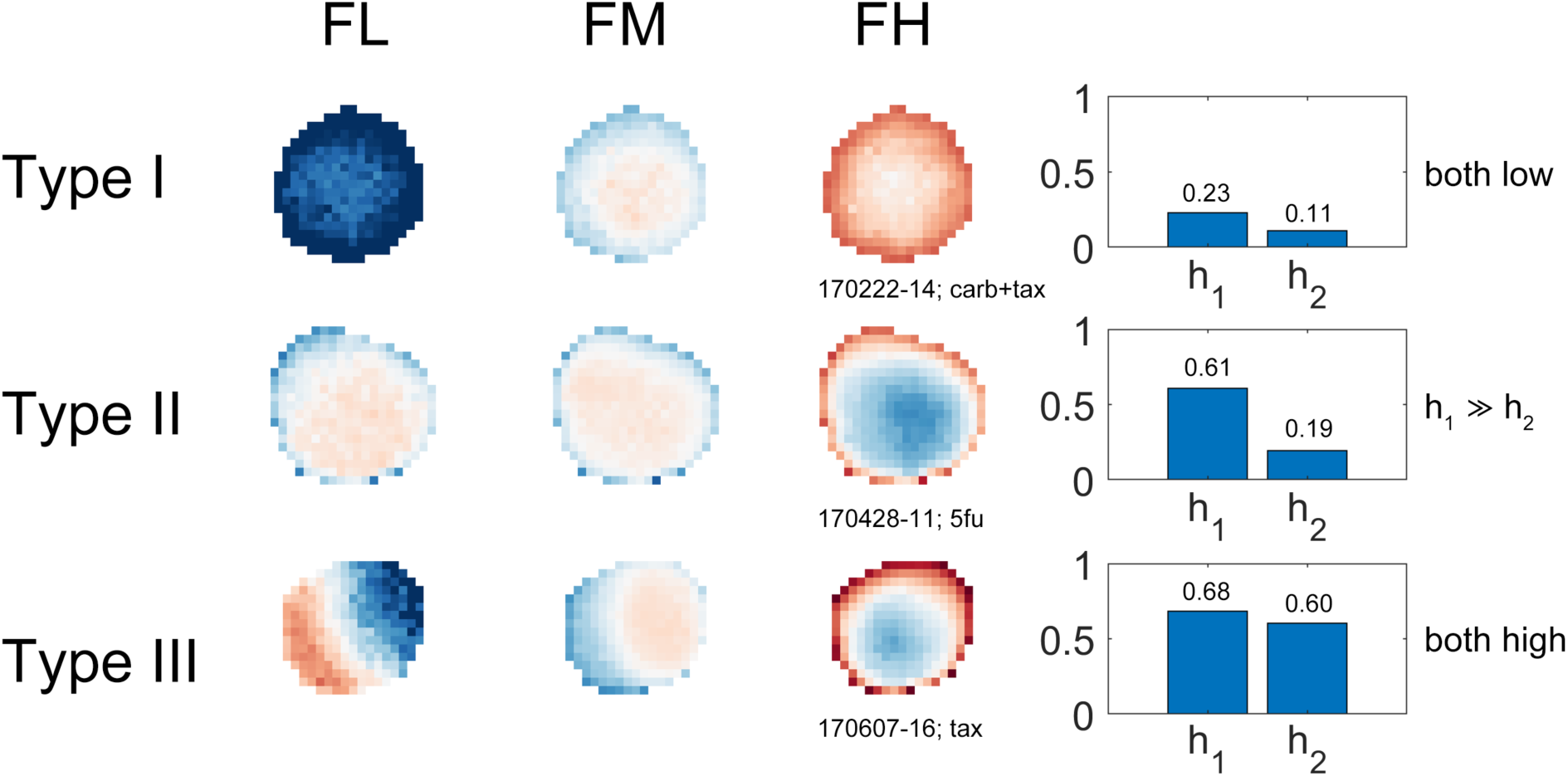
Three types of samples that have different levels of same-mask heterogeneity and cross-mask heterogeneity, with scores on the right.

Time-lapse images offer an additional layer of understanding of the spatial evolution of drug effects or sample conditions. In Fig. 7 for sample “170317-9” treated with nocodazole, the blue response pattern (mid-frequency suppression and low- and high-frequency enhancement) grows stronger over time before saturation, which indicates that nocodazole’s suppression of microtubule polymerization begins at the outer periphery and slowly penetrates the core of the sample. As another example, the red area in the TDS image of sample “170606-15” becomes stronger until around 9 hours, when the sample displays an overall suppression across the entire sample.

**Fig. 7.**
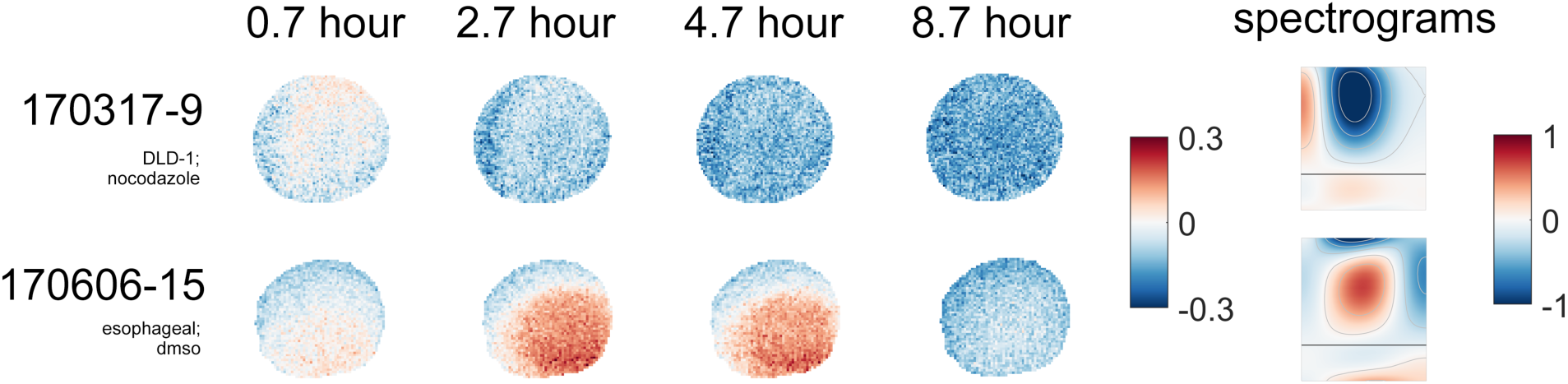
Time-lapse TDS image of samples responding to drugs. Sample “170317-9”: Response of a DLD spheroid sample treated with 10 μM nocodazole (same as in Fig. 3d) with the G2 mask, showing a silent core shortly after drug was added <u>(0.7 hours)</u>, which was “invaded” by the drug and later achieved a spatially homogeneous response <u>(2.7 hours to 8.7 hours)</u>. Sample “170606-15”: Response of an esophageal biopsy sample in the control medium, also under the G2 mask.

## 4 Comparison with inverted Selective-Plane Illumination Microscopy

Given that BDI is a 3D imaging technology that uses low-coherence light, the different drug response phenotypes revealed by TDSI may be related to different types of tissues in a certain region of a sample. A complementary 3D imaging technique is iSPIM that produces microscopic images of 3D slices with high lateral and axial resolutions, which allows us to distinguish features in the images. Therefore, by comparing TDSI to iSPIM, we can investigate whether the heterogeneity related to drug response variability from TDSI is also present in microscopic images, i.e. link dynamic information from functional imaging with the histology of biological tissues.

As an example, TDSI maps for sample 190801-15 are compared against its iSPIM images and H&E histology images in Fig. 8. This sample is a pinch biopsy from a patient who was resistant to neoadjuvant therapy. The biodynamic response is mapped for the tissue response to 5-fluorouracil. In the TDSI map, there is a distinct central region (purple) contrasted with the outer regions (orange). In the bivariate colormap, purple represents broadband excitation with a blue shift, indicating an activated response of the tissue to the 5-fu. The central region is likely to be naturally hypoxic, which can affect the mechanism of the drug. In a related study of esophageal cancer patients, a blue shift induced by 5-fu is representative of a beneficial response to the chemotherapy. In contrast, in the peripheral tissue, that is hyperoxic, the broadband excitation is accompanied by a red shift. Furthermore, the green region indicates overall suppressed activity (broad inhibition accompanied with a red shift of slower intracellular speeds). In comparison to the TDSI, in the iSPIM image the lower part matches the lower part of the histology image containing a large concentration of DNA. The upper part of the iSPIM image is cytoplasm or unstained tissues, matching the lack of nuclei in the upper part of the histology image, which potentially indicates collagenous connective tissues. Although the image orientations were not registered between the techniques, the images in Fig. 8 demonstrate spatial polarity in the tissue types. Future studies to compare TDSI to iSPIM would register sample orientation to identify TDS spectral signatures of connective tissue relative to epithelial tissue. The work presented here is proof-of-principle that spatial heterogeneity can be observed in both imaging modalities.

**Fig. 8.**
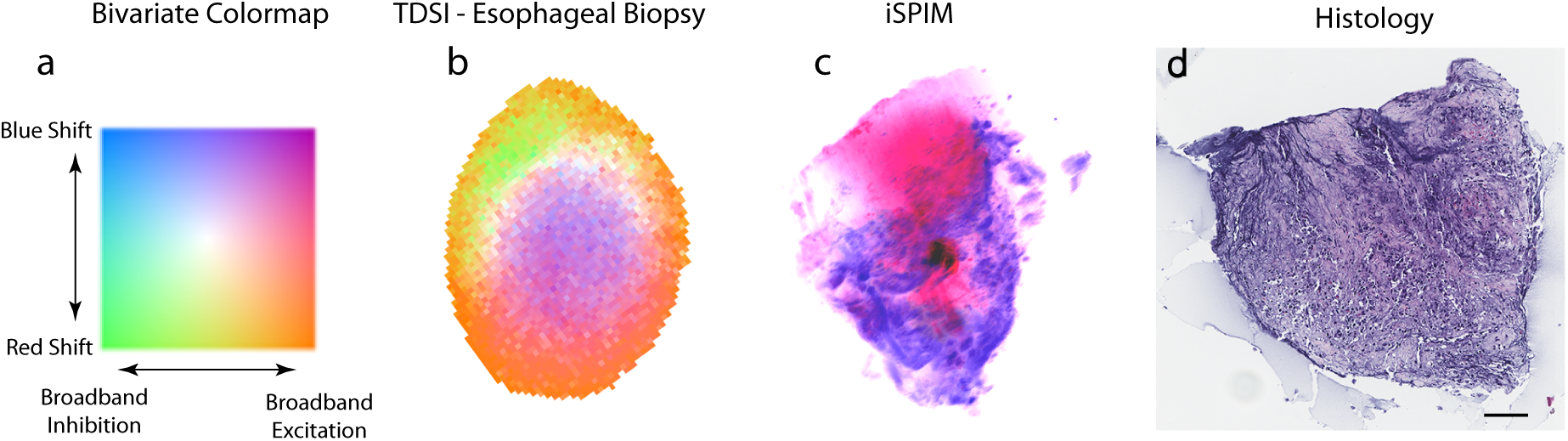
TDSI map, iSPIM 3D-reconstructed image and H&E histology image for an esophageal biopsy sample. All scale bars are 100 μm. a) bivariate colormap representing G0 (broadband inhibition) and G1 (blue shift). b) bivariate TDSI map of human esophageal biopsy responding to 5-fluorouracil (5-fu) with G0 and G2 as the two masks. The color map and color scale are the same as in Fig. 3. c) the iSPIM image with DRAQ 5 (blue) and Eosin (pink) for the same sample in a), and d) Histology image. The orientations of the images are not registered.

In this comparison, the advantage of iSPIM is the ability to acquire microscopic image with high resolution and detailed cellular-level structural information with acceptable speed. This is in contrast to TDSI which achieves only tissue-scale imaging resolution (approximately 15 micron voxel size). On the other hand, TDSI is a functional imaging method that is highly specific to the action of drugs on tissue dynamics, while iSPIM can only provide structural information without being specific to drug response. Furthermore, TDSI is nondestructive, allowing longitudinal studies that can span several days, including the ability to apply drugs and to clear them. Both techniques have the advantage of imaging into three-dimensional tissues, which is a key aspect of maintaining the tumor microenvironment during imaging.

## 5 Discussion

Biodynamic imaging is a tool that is sensitive to intracellular dynamics and has been applied successfully to phenotypic profiling to predict patient outcomes. For instance, sample motility and dynamic biomarkers have been shown to be consistent and reliable indicators of pharmacodynamics effects. However, these biomarkers are usually calculated as whole-sample averages when used in classification and similarity analyses, overlooking intra-sample heterogeneity. The introduction of TDSI soles this problem by evaluating the responses of subregions of the sample to reveal new information on the complex spatial structures in sample drug response, which is supported by evidence from other imaging techniques like iSPIM and histology. Visualization of sample heterogeneity is facilitated with a bivariate color representation and is quantitatively characterized by *h*_1_ and *h*_2_ scores. This imaging method is further extended to generate whole-sample time-lapse TDSI maps, providing a novel method to monitor drug mechanisms.

In addition to visualizing sample heterogeneity, TDSI maps can provide additional information and improve classification accuracy when evaluating anti-cancer drug effectiveness on a patient level. For samples with regions of opposite responses, the whole-sample average spectrogram may suggest a mild response to the drug, making the sample and the patient appear to be less sensitive to the treatment. The proposed solution here is to introduce additional biomarkers that characterize regional drug responses. As an example, the sample shown in Fig. 3b can be split into two regions based on the sign of G1 biomarker values (which can be related to the sample’s heterogeneous structure), and the feature values of these two regions can be calculated. The set of feature values that capture both sample average response as well as regional response would provide a more comprehensive assessment of the patient.

TDSI can be extended for further imaging and analysis applications. Since BDI is a 3D imaging technique and achieves depth selection with coherence gating, a volumetric TDS image can be generated by scanning different slices of a sample. Also, time-lapse TDS analysis is a quantitative approach to visualize drug-action time dependence. Features like delay and distribution related to pharmacokinetics and pharmacodynamics (PK/PD) could be obtained from time-lapse images to provide insight into processes like dose-response relationships.

Challenges to TDSI include sample immobilization and multiple light scattering. TDSI evaluates the drug spectral response on the pixel level and requires that the same part of the sample is imaged throughout the experiment. This requires the sample to maintain the same lateral and axial positions. In addition, multiple light scattering induces aberrations of the image and reduces the signal-to-noise ratio, which makes TDSI more effective at shallower depths.

TDSI is an important extension to the current suite of BDI modalities (OCI, MCI, TDS). MCI maps are simple and intuitive functional images that visualize sample motility and have revealed the contrast between a viable shell and a necrotic core for rat tumor spheroids(27). TDSI, by comparison, generates a set of more detailed functional maps that complement MCI because the critical frequencies in spectral masks used in TDSI are related to specific types of intracellular components and motions, offering a comprehensive view of changes occurring in the sample. TDSI is a versatile functional imaging method that could provide new information for drug-response profiling and has the potential for improving predictions of response to therapy and drug screening.

## Acknowledgements

This work was supported by grants NSF 1911357-CBET and NIH NIBIB 1RO1EB016582 and by the Purdue University Cancer Center.

## Biographies

**Zhe Li** received his PhD in physics from Purdue University in 2020. He studied intracellular dynamics of living tissues using Doppler fluctuation spectroscopy under the direction of Dr. David Nolte. He currently works on model verification and validation products at MathWorks, Natick, Massachusetts, USA.

**Guang Li** is a Ph.D candidate in biomedical engineering department of Tulane University. He received his BS in Dalian University of Technology in China and MS in Tulane. Working in Translational Biphotonics Laboratory directed by Dr. J. Q. Brown, he is focusing on image processing of dual-view inverted Selective Plane Illumination Microscopy (diSPIM) and improvement of tissue clearing and fluorescent staining protocols. Besides, he also aims to apply light sheet microscopy to other practical areas.

**Sharon E. Fox, MD, PhD**, is the Associate Director of Research and Development in the Department of Pathology at LSU Health Sciences Center, New Orleans, and an anatomic pathologist at the Southeast Louisiana Veterans Healthcare System. She received her MD from Harvard Medical School in 2008, and PhD from MIT in 2012. Her research interests include digital pathology, autopsy pathology, and the interaction between healthcare providers and image data.

**Shadia I Jalal** MD, is an Associate Professor of Medicine in the Department of Medicine-Division of Hematology Oncology at Indiana University School of Medicine. She is a thoracic oncologist and Phase I physician. She is a clinical researcher that designs, runs and serves as the principal investigator of many clinical trials. Her research focus is in lung and esophageal cancers.

**Bihe Hu** received her BS and MS in Biomedical Engineering from Huazhong University of Science and Technology, where her research interest was in the area of 3D whole-brain imaging at single-cell resolution. She received her Ph.D. from Tulane University in 2020. During her Ph.D. study, her research focused on the development and optimization of light sheet fluorescence microscopy and its application in 3D histopathology.

**J. Quincy Brown** is the Paul H. and Donna D. Flower assistant professor in the Department of Biomedical Engineering at Tulane University, and a program member in the Tulane Cancer Center. His research interests include point-of-care digital pathology for cancer diagnosis. He is a member of SPIE, OSA, IEEE-EMBS, and BMES.

**John Turek** received his PhD from the University of Illinois at Urbana in Veterinary Medical Science. He is a professor in the Department of Basic Medical Sciences at Purdue University College of Veterinary Medicine. For the past 16 years, his research has been in the area of biodynamic imaging with a focus on the use of 3-D cell cultures and BDI for derisking drug discovery and for cancer chemotherapy selection.

**David D. Nolte** is the E.M Purcell distinguished professor of physics at Purdue University. He received his BS degree from Cornell University in 1981 and his PhD from the University of California at Berkeley in 1988, followed by a postdoctoral appointment at AT&T Bell Labs. He has been elected as a fellow of the Optical Society of America, a fellow of the American Physical Society, and a fellow of the AAAS. He is the technical founder of two biotech startup companies.

## References

1. M. Gerlinger et al., “Intratumor Heterogeneity and Branched Evolution Revealed by Multiregion Sequencing,” New England Journal of Medicine 366(10), 883–892 (2012).

2. A. S. Little, P. D. Smith, and S. J. Cook, “Mechanisms of acquired resistance to ERK1/2 pathway inhibitors,” Oncogene 32(10), 1207–1215 (2013).

3. D. Barbone et al., “Mammalian target of rapamycin contributes to the acquired apoptotic resistance of human mesothelioma multicellular spheroids,” Journal Of Biological Chemistry 283(19), 13021–13030 (2008).

4. G. A. Cirkel et al., “Tumor heterogeneity and personalized cancer medicine: are we being outnumbered?,” Future Oncology 10(3), 417–428 (2014).

5. O. Tredan et al., “Drug resistance and the solid tumor microenvironment,” Journal Of The National Cancer Institute 99(19), 1441–1454 (2007).

6. A. Tomida, and T. Tsuruo, “Drug resistance mediated by cellular stress response to the microenvironment of solid tumors,” Anti-Cancer Drug Design 14(2), 169–177 (1999).

7. M. Esteller, and J. G. Herman, “Cancer as an epigenetic disease: DNA methylation and chromatin alterations in human tumours,” Journal Of Pathology 196(1), 1–7 (2002).

8. P. A. Kenny, and M. J. Bissell, “Tumor reversion: Correction of malignant behavior by microenvironmental cues,” International Journal of Cancer 107(5), 688–695 (2003).

9. X. Xu, M. C. Farach-Carson, and X. Q. Jia, “Three-dimensional in vitro tumor models for cancer research and drug evaluation,” Biotechnology Advances 32(7), 1256–1268 (2014).

10. P. D. Caie et al., “High-Content Phenotypic Profiling of Drug Response Signatures across Distinct Cancer Cells,” Molecular Cancer Therapeutics 9(6), 1913–1926 (2010).

11. H. S. Seol et al., “A patient-derived xenograft mouse model generated from primary cultured cells recapitulates patient tumors phenotypically and genetically,” Journal Of Cancer Research And Clinical Oncology 139(9), 1471–1480 (2013).

12. H. Y. Deng, C. M. Wang, and Y. Fang, “Label-free cell phenotypic assessment of the molecular mechanism of action of epidermal growth factor receptor inhibitors,” Rsc Advances 3(26), 10370–10378 (2013).

13. P. Yu et al., “Holographic optical coherence imaging of rat osteogenic sarcoma tumor spheroids,” Applied Optics 43(25), 4862–4873 (2004).

14. P. Yu et al., “Holographic optical coherence imaging of tumor spheroids,” Appl Phys Lett 83(3), 575–577 (2003).

15. E. Cuche, F. Bevilacqua, and C. Depeursinge, “Digital holography for quantitative phase-contrast imaging,” Optics Letters 24(5), 291–293 (1999).

16. P. Marquet et al., “Digital holographic microscopy: a noninvasive contrast imaging technique allowing quantitative visualization of living cells with subwavelength axial accuracy,” Optics Letters 30(5), 468–470 (2005).

17. N. T. Shaked et al., “Whole-cell-analysis of live cardiomyocytes using wide-field interferometric phase microscopy,” Biomedical Optics Express 1(2), 706–719 (2010).

18. E. Beaurepaire et al., “Full-field optical coherence microscopy,” Opt. Lett. 23(4), 244–246 (1998).

19. Z. Yaqoob et al., “Homodyne en face optical coherence tomography,” Optics Letters 31(12), 1815–1817 (2006).

20. K. Jeong, J. J. Turek, and D. D. Nolte, “Imaging Motility Contrast in Digital Holography of Tissue Response to Cytoskeletal Anti-cancer Drugs,,” Optics Express 15(14057–14064 (2007).

21. D. D. Nolte et al., “Holographic tissue dynamics spectroscopy,” Journal of Biomedical Optics 16(8), 087004–087013 (2011).

22. D. D. Nolte et al., “Tissue dynamics spectroscopy for phenotypic profiling of drug effects in three-dimensional culture,” Biomed. Opt. Express 3(11), 2825–2841 (2012).

23. D. Merrill et al., “Intracellular Doppler Signatures of Platinum Sensitivity Captured by Biodynamic Profiling in Ovarian Xenografts,” Nature Scientific Reports 6(18821 (2016).

24. M. R. Custead et al., “Predictive value of ex vivo biodynamic imaging in determining response to chemotherapy in dogs with spontaneous non-Hodgkin’s lymphomas: a preliminary study,” Convergent Science Physical Oncology 1(1), 015003 (2015).

25. R. An et al., “Biodynamic imaging of live porcine oocytes, zygotes and blastocysts for viability assessment in assisted reproductive technologies,” Biomedical Optics Express 6(3), 963–976 (2015).

26. D. D. Nolte et al., “Holographic tissue dynamics spectroscopy,” J. Biomed. Opt. 16(8), 087004-087004-087013 (2011).

27. K. Jeong, J. J. Turek, and D. D. Nolte, “Volumetric motility-contrast imaging of tissue response to cytoskeletal anti-cancer drugs,” Opt. Express 15(21), 14057–14064 (2007).

28. C. Apelian et al., “Dynamic full field optical coherence tomography: subcellular metabolic contrast revealed in tissues by interferometric signals temporal analysis,” Biomed Opt Express 7(4), 1511–1524 (2016).

29. W. Tan et al., “Optical coherence tomography of cell dynamics in three-dimensional tissue models,” Optics Express 14(16), 7159–7171 (2006).

30. C. Joo et al., “Diffusive and directional intracellular dynamics measured by field-based dynamic light scattering,” Optics Express 18(3), 2858–2871 (2010).

31. G. Farhat et al., “Detecting apoptosis using dynamic light scattering with optical coherence tomography,” Journal of Biomedical Optics 16(7), 070505 (2011).

32. J. Lee et al., “Dynamic light scattering optical coherence tomography,” Optics Express 20(20), 22262–22277 (2012).

33. A. L. Oldenburg et al., “Inverse-power-law behavior of cellular motility reveals stromal-epithelial cell interactions in 3D co-culture by OCT fluctuation spectroscopy,” Optica 2(10), 877–885 (2015).

34. N. J. J. Arezza, M. Razani, and M. C. Kolios, “Dynamic light scattering optical coherence tomography to probe motion of subcellular scatterers,” J Biomed Opt 24(2), 1–7 (2019).

35. H. Sun, “Dynamic Holography in Semiconductors and Biomedical Optics,” Purdue University, West Lafayette, IN, USA (2016).

36. Z. Li et al., “Doppler fluctuation spectroscopy of intracellular dynamics in living tissue,” J. Opt. Soc. Am. A 36(4), 665–677 (2019).

37. M. Harrower, and C. A. Brewer, “ColorBrewer.org: An Online Tool for Selecting Colour Schemes for Maps,” The Cartographic Journal 40(1), 27–37 (2003).

38. A. J. Teuling, R. Stöckli, and S. I. Seneviratne, “Bivariate colour maps for visualizing climate data,” International Journal of Climatology 31(9), 1408–1412 (2011).

39. K. G. Baum et al., “Techniques for Fusion of Multimodal Images: Application to Breast Imaging,” 2006 International Conference on Image Processing 2521–2524 (2006).

40. J. Bernard et al., “A survey and task-based quality assessment of static 2D colormaps,” SPIE/IS&T Electronic Imaging 9397((2015).

41. S. Bremm et al., “Assisted Descriptor Selection Based on Visual Comparative Data Analysis,” Computer Graphics Forum 30(3), 891–900 (2011).

42. M. Steiger et al., “Explorative Analysis of 2D Color Maps,” (2015).

43. M. Steiger et al., “Visual Analysis of Time-Series Similarities for Anomaly Detection in Sensor Networks,” Computer Graphics Forum 33(3), 401–410 (2014).

44. R. Bhatia, and C. Davis, “A Better Bound on the Variance,” The American Mathematical Monthly 107(4), 353–357 (2000).

45. B. Hu, G. Li, and J. Q. Brown, “Enhanced resolution 3D digital cytology and pathology with dual-view inverted selective plane illumination microscopy,” Biomed Opt Express 10(8), 3833–3846 (2019).

46. B. Hu, D. Bolus, and J. Q. Brown, “Improved contrast in inverted selective plane illumination microscopy of thick tissues using confocal detection and structured illumination,” Biomed Opt Express 8(12), 5546–5559 (2017).

47. G. Li, B. Hu, and J. Q. Brown, “An approach of 3D reconstruction for images by Dual-view Inverted Selective Plane Illumination Microscopy (diSPIM),” in The Optical Society, Biophotonics Congress: Optics in the Life Sciences Congress 2019 (BODA,BRAIN,NTM,OMA,OMP) NW5C.5 (2019).

48. S. Preibisch, S. Saalfeld, and P. Tomancak, “Globally optimal stitching of tiled 3D microscopic image acquisitions,” Bioinformatics 25(11), 1463–1465 (2009).

49. M. G. Giacomelli et al., “Virtual Hematoxylin and Eosin Transillumination Microscopy Using Epi-Fluorescence Imaging,” PLOS ONE 11(8), e0159337 (2016).

50. H. Peng et al., “V3D enables real-time 3D visualization and quantitative analysis of large-scale biological image data sets,” Nature biotechnology 28(4), 348–353 (2010).

51. H. Peng et al., “Extensible visualization and analysis for multidimensional images using Vaa3D,” Nature protocols 9(1), 193–208 (2014).

