## Supplemental Information for "Tissue Dynamics Spectroscopic Imaging: Functional Imaging of Heterogeneous Cancer Tissue"

### S.1 Condensed Data Format

An intermediate data processing step generates a condensed data format (CDF) in files designed for purposes like TDSI, introduced to allow quick calculation of the normalized spectrum of a randomly shaped area while using only a small amount of additional storage. This is achieved by using spatial down-sampling followed by frequency down-sampling. A comparison of the formats for data needed to reconstruct a single-loop for a single-well spectrum is shown in Table 1.

**Table 1** Comparison of three data formats for speckle fluctuation spectrum reconstruction

| **Type** | **Format** | **Disk Space** | **Notes** |
|---|---|---|---|
| Raw Images | 2048 × 800 × 800 × 16bit uint16 | 2.44 GB | 16bit TIFF files |
| Analyzed Data | 2048 × 256 × 256 × 64bit double | 1 GB | All reconstructed frames |
| Condensed Data format | 130 × 128 × 128 × 64bit double +<br>1 × 128 × 128 × 64bit double | 16 MB | Down-sampled spectrum and normalization factor |

Spatial down-sampling (Fig. S1a, b) enables the analysis of spectral features with low noise at good spatial resolution. The spectrum of a single sample can be too noisy to extract spectral features by methods like curve fitting. Therefore, each sample is divided into 2 × 2 pixel squares (referred to as "TDSI pixels" below) as the basic unit for spectral averaging. The point-spread function diameter on the reconstructed imagine plane is approximately 15 μm spanned by three pixels. Therefore, the 2x2 binning causes only a small decrease in spatial resolution and averaging over the four pixels represents a coherent sum (within the spatial coherence area). Because biodynamic imaging is an interferometric technique using short coherence, each image is reconstructed from backscattered light from a selected depth in the sample. Therefore, each TDSI pixel represents a voxel of the sample. As a nominal choice that balances spatial resolution and spectral resolution, the spectrum of each TDSI pixel is the average over the 4 pixels, given by

$$S_{\text{raw}}\left(\tilde{i},\tilde{j};f\right)=\frac{1}{4}\sum_{(i,j)\in\sigma(\tilde{i},\tilde{j})}S_{\text{raw}}\left(i,j;f\right) \tag{1}$$

where $\sigma(\tilde{i},\tilde{j})$ is the area $\{2\tilde{i}-1,2\tilde{i}\}\times\{2\tilde{j}-1,2\tilde{j}\}$ in the original image, and $(\tilde{i},\tilde{j})$ are the coordinates in the sub-sampled image.

Frequency down-sampling reduces the frequency components stored while still allowing high-precision reconstruction. Our data acquisition format (discussed in Section 2.2) generates spectra of 1024 frequency components evenly spaced on a linear scale. However, TDS analysis is based on spectra on a log-log scale, which have sparse points in the low frequencies and dense points in high frequencies. Our frequency down-sampling method remaps the 1024 components to 130 that are evenly spaced on the log scale. The low-frequency part is up-sampled with interpolation, while the high frequency uses a binning method and preserves the average. The original normalization factor $M(\tilde{i},\tilde{j})=\sum_f S_{\mathrm{raw}}(\tilde{i},\tilde{j};f)$ is stored to maintain the same normalization. In our data analysis, the order of averaging is not relevant

$$\sum_f \sum_{(\tilde{i},\tilde{j})\in\sigma} S(\tilde{i},\tilde{j};f)=\sum_{(\tilde{i},\tilde{j})\in\sigma}\sum_f S(\tilde{i},\tilde{j};f) \tag{2}$$

and the normalized spectrum is reconstructed by

$$S_{\sigma,\mathrm{norm}}(f)=\frac{S_{\sigma,\mathrm{raw}}(f)}{\sum_f S_{\sigma,\mathrm{raw}}(f)}=\frac{\sum_{(\tilde{i},\tilde{j})\in\sigma} S(\tilde{i},\tilde{j};f)}{\sum_f \sum_{(\tilde{i},\tilde{j})\in\sigma} S(\tilde{i},\tilde{j};f)}=\frac{\sum_{(\tilde{i},\tilde{j})\in\sigma} S(\tilde{i},\tilde{j};f)}{\sum_{(\tilde{i},\tilde{j})\in\sigma}\sum_f S(\tilde{i},\tilde{j};f)}=\frac{\sum_{(\tilde{i},\tilde{j})\in\sigma} S(\tilde{i},\tilde{j};f)}{\sum_{(\tilde{i},\tilde{j})\in\sigma} M(\tilde{i},\tilde{j})} \tag{3}$$

Eq. (3) shows that once the TDSI pixel-wise down-sampled spectra $S(\tilde{i},\tilde{j};f)$ and normalization factors $M(\tilde{i},\tilde{j})$ are obtained, the normalized spectrum of any given region $\sigma$ can be reconstructed. On log-log plots, the difference between spectrograms reconstructed from CDF files and those reconstructed from original pixels is negligible. The comparisons are presented in Fig. S1c and 3d.

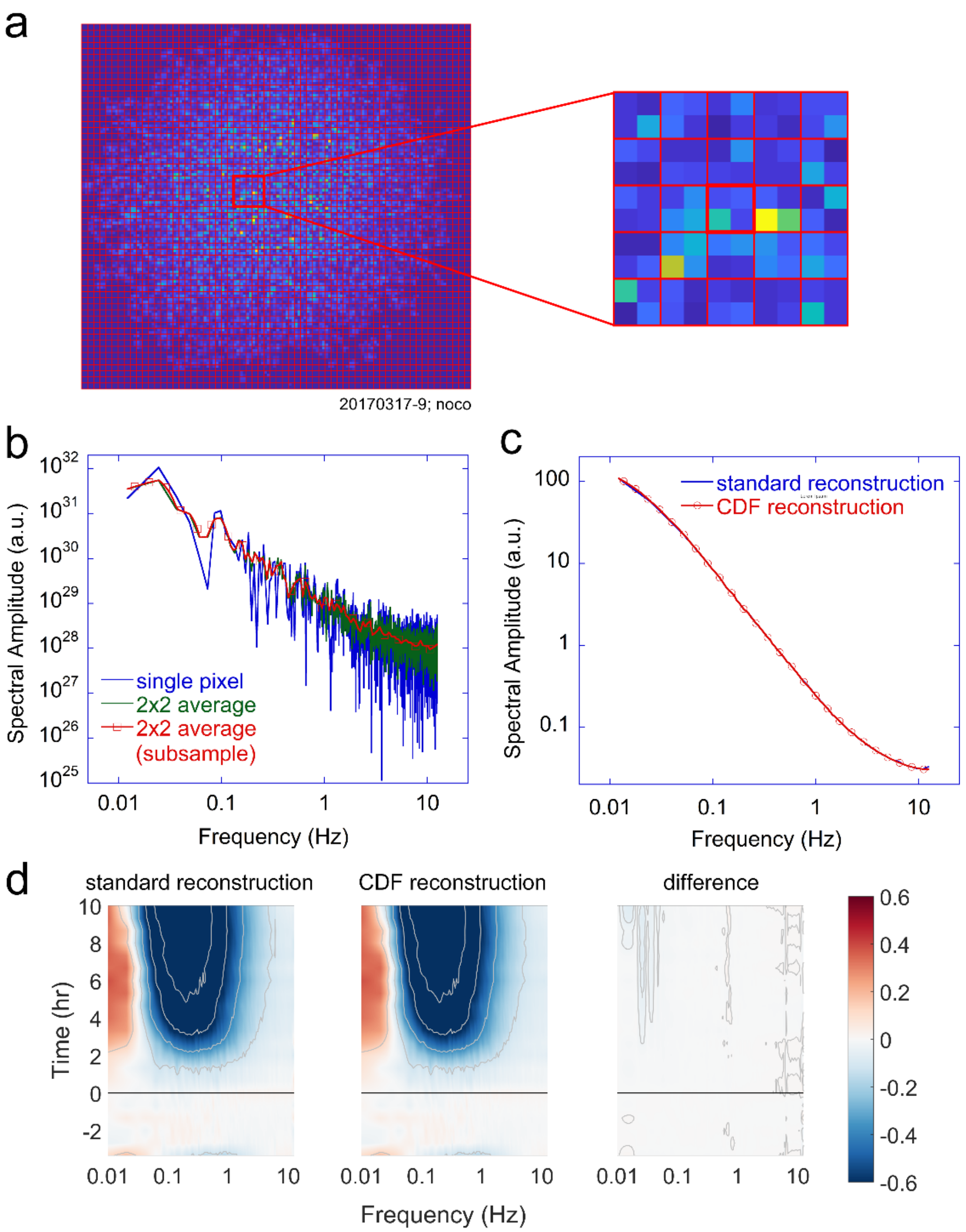


Fig. S1 Demonstration of the condensed data format (CDF). The sample used in this figure is a DLD-1 spheroid treated with 10 μM nocodazole. a) Spatial down-sampling, with a 10 × 10 pixel region enlarged. b) Frequency down-sampling, with the spectrum of a single pixel in the 2 × 2 px area (labeled in Fig. S1a with heavy border lines), the average spectrum of 4 pixels in the area, and the down-sampled spectrum with 130 components evenly spaced on the log frequency axis. c) Comparison of a sample average spectrum reconstructed from raw data and from the CDF. d) Whole-sample full-length spectrogram reconstructed from raw data and from a CDF file. The relative difference is <0.5%. Nocodazole was added at $t = 0$.
